# PTM-dCN: Latent Space Control for Post-Translational Modification–Aware Protein Design

**DOI:** 10.64898/2026.05.06.714367

**Authors:** Sitao Zhang, Rui Qing, Tianming Huang, Edward Chen

## Abstract

Post-translational modifications (PTMs) are critical for protein function, yet their precise design by harnessing site specific information derived from native proteins remains challenging. Here, we present a deep learning–based PTM design framework that integrates latent diffusion models with ControlNet for sequence generation with site-specific PTM-control. The framework incorporates a PTM-aware protein language model featuring extractor, trained on a curated SwissProt PTM dataset with specialized modification tokens. Through de novo generation of protein sequences with designated PTM sites, our framework facilitates the exploration of PTM-driven functional landscapes and advances position-aware protein engineering.

## 1 Introduction

Post-translational modifications (PTMs) play a critical role in protein functional diversity, regulating activity, stability, localization, and signaling processes Lee et al. (2023); Knorre et al. (2009). Despite their importance, most computational design methods focus on canonical amino-acid sequences and do not explicitly model or generate PTM-containing proteins. In contrast, existing PTM-focused approaches are primarily limited to prediction models, which only identify potential PTM sites and predict their impact on protein function, but do not necessarily directly design sequences with PTM modifications Shrestha et al. (2024); Wang et al. (2020); Peng (2024). Traditional rational design strategies can also introduce PTMs only based on fixed consensus motifs, but fail to consider the broader sequence context, especially motifs conformationally close but far apart in the primary sequence, limiting their generalizability Kholodenko & Okada (2021). Together, these gaps underscore the lack of methods capable of context-aware PTM sequence generation in full protein level.

Generating novel protein sequences with precise PTM is challenging because it requires redesigning local sequence contexts around modification sites that can be built into global coherence. The difficulty increases when multiple PTM types or sites need to be inserted simultaneously, necessitating models that can reconcile local constraints with the plausibility of the whole sequence.

To address this, we introduce PTM-dCN, a diffusion-based framework for protein sequence generation with explicit, site-specific PTM design. Inspired by ControlNet Zhang et al. (2023), which achieves state-of-the-art performance in fine-grained, structure-conditioned image generation. We adapt this paradigm to protein sequences via conditioning the generative process on predefined PTM sites of different types. PTM-dCN redesigns local amino-acid contexts without disrupting the overall sequence and structure. We also constructed a high-quality PTM-annotated dataset derived from SwissProt, and built a vocabulary with specialized PTM tokens to explicitly represent and annotate modified residues.

We evaluated PTM-dCN on lactate dehydrogenase (LDH) or isocitrate dehydrogenase (IDH) datasets. The model generates novel PTM-containing sequences with high predicted foldability, which preserves or enhances PTM occurrence probabilities at target sites, and adapts PTM motifs in a sequence-context–dependent manner. This is the first framework for PTM-aware protein sequence engineering, providing a generalizable approach for controllable protein design with precise PTM insertion, which enables a novel methodology for functional tuning.

## 2 Experiments and Results

### 2.1 Overview of PTM-dCN

PTM-dCN enables precise site-specific design protein sequence of a diffusion-based framework. The model consists of a latent diffusion backbone and an auxiliary ControlNet module for fine-grained conditioning (Figure 1).

**Figure 1:**
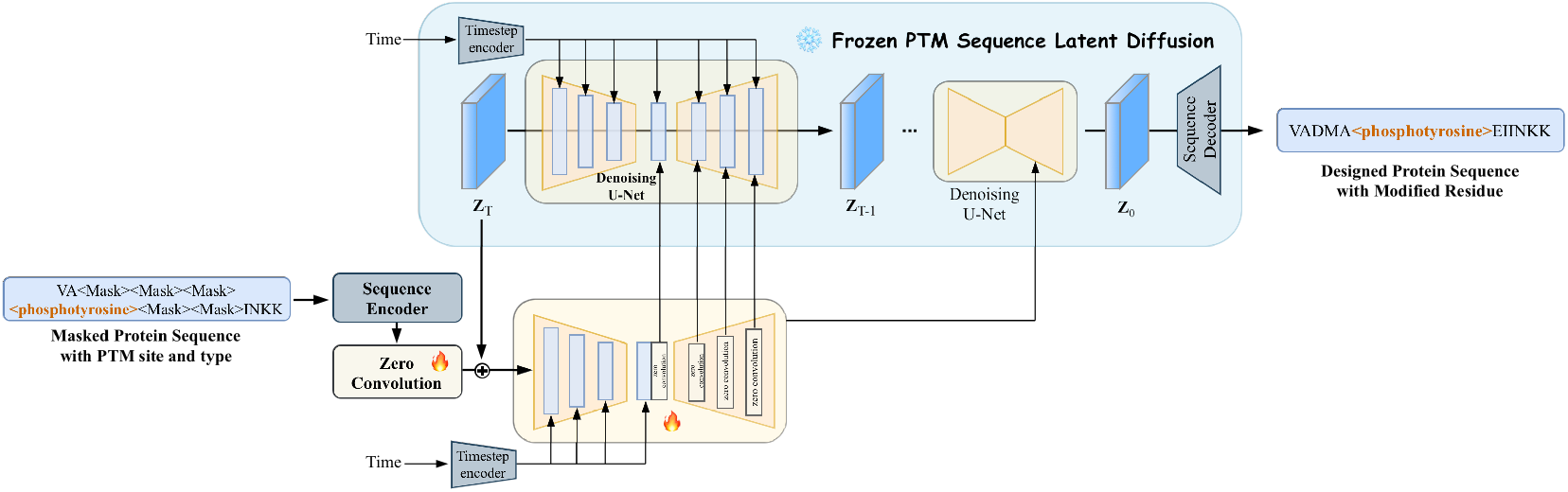
The architecture of PTM-dCN.

The latent diffusion backbone is built on our previous work, PRO-LDM Zhang et al. (2025), and is extended with an expanded token vocabulary to support the learning and generation of complete protein sequences containing modified residues with PTM information. This design allows the model to capture global sequence distributions while explicitly representing PTMs at the token level.

Inspired by fine-grained structure-conditioned image generation using ControlNet Zhang et al. (2023), we introduce a ControlNet module in parallel to the diffusion backbone to impose explicit constraints on PTM sites and the underlying sequence design scaffold. By conditioning the diffusion process on site-specific PTM information, the model constructs the amino acid context surrounding modification sites at the global level, enabling precise and controllable PTM-aware sequence design without compromising the global generative capacity at the backbone model.

### 2.2 PTM-dCN Generates Highly Foldable Protein Sequences with PTMs

The staged training strategy is adopted to ensure both high-quality sequence representations and controllable generation. Specifically, (i) the autoencoder (AE) module was first trained independently to obtain robust PTM sequence embeddings; (ii) the diffusion backbone was then trained to endow the model with strong protein sequence generation capability; and (iii) after convergence of the backbone, its parameters were frozen and the ControlNet module was trained separately for efficient and stable conditioning on PTM sites.

During AE training, sequence recovery accuracy is used as the primary evaluation metric. As shown in Figure A.1, on the SwissProt-PTM dataset, the AE achieves around 50% sequence recovery, with the corresponding perplexity reduced to 8.16. The impact of encoder capacity and latent dimensionality is further examined by employing ESM2 encoders of different parameter scales and varying latent dimensions. Under matched settings, increasing encoder capacity and latent dimensionality leads to improved sequence recovery, with sequence recovery increasing from 0.62 to 0.55 and perplexity decreasing from 8.78 to 8.16 when changing an 8M ESM2 encoder with latent dimension 64 to a 35M encoder with latent dimension 256. This trend indicates that larger encoder and higher dimension consistently improve sequence recovery performance.

After AE training, embeddings of ten randomly selected protein sequences are projected into a two-dimensional space for visualization. For each sequence, both the canonical amino-acid version and the corresponding PTM-containing version are mapped. Figure 2 shows that embeddings of the two versions of the same sequence are closely clustered but not fully overlapping. This behavior suggests that the learned latent space preserves overall sequence identity while explicitly distinguishing modified residues, enabling discrimination between canonical and PTM-containing sequences at the representation level.

**Figure 2:**
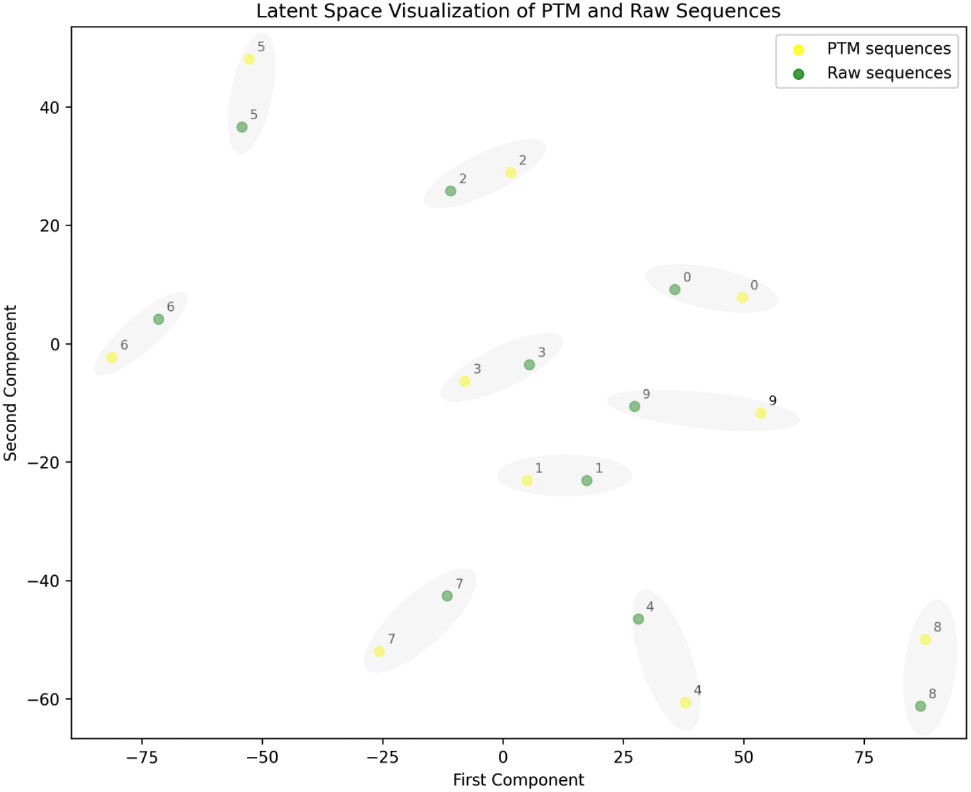
Latent Space Visualization of Canonical vs PTM-Containing Protein Sequences.

Following embedding analysis, we subsequently investigated PTM-dCN’s ability to generate PTM-containing sequences within specific functional contexts by fine-tuning the diffusion backbone on lactate dehydrogenase (EC 1.1.1.27) or isocitrate dehydrogenase (EC 1.1.1.42) datasets. After training the AE and diffusion modules separately on the LDH and IDH datasets, the model achieved over 85% sequence reconstruction accuracy and was able to generate novel PTM-containing protein sequences with predicted pLDDT *>*90. Furthermore, by using AlphaFold3 to specifically annotate PTM sites and types, we performed structure predictions on the same sequences both with and without PTM annotations. The results showed that sequences with annotated PTM sites consistently exhibited higher predicted pLDDT scores, indicating enhanced foldability compared to their unmodified counterparts (Figure 3 ).

**Figure 3:**
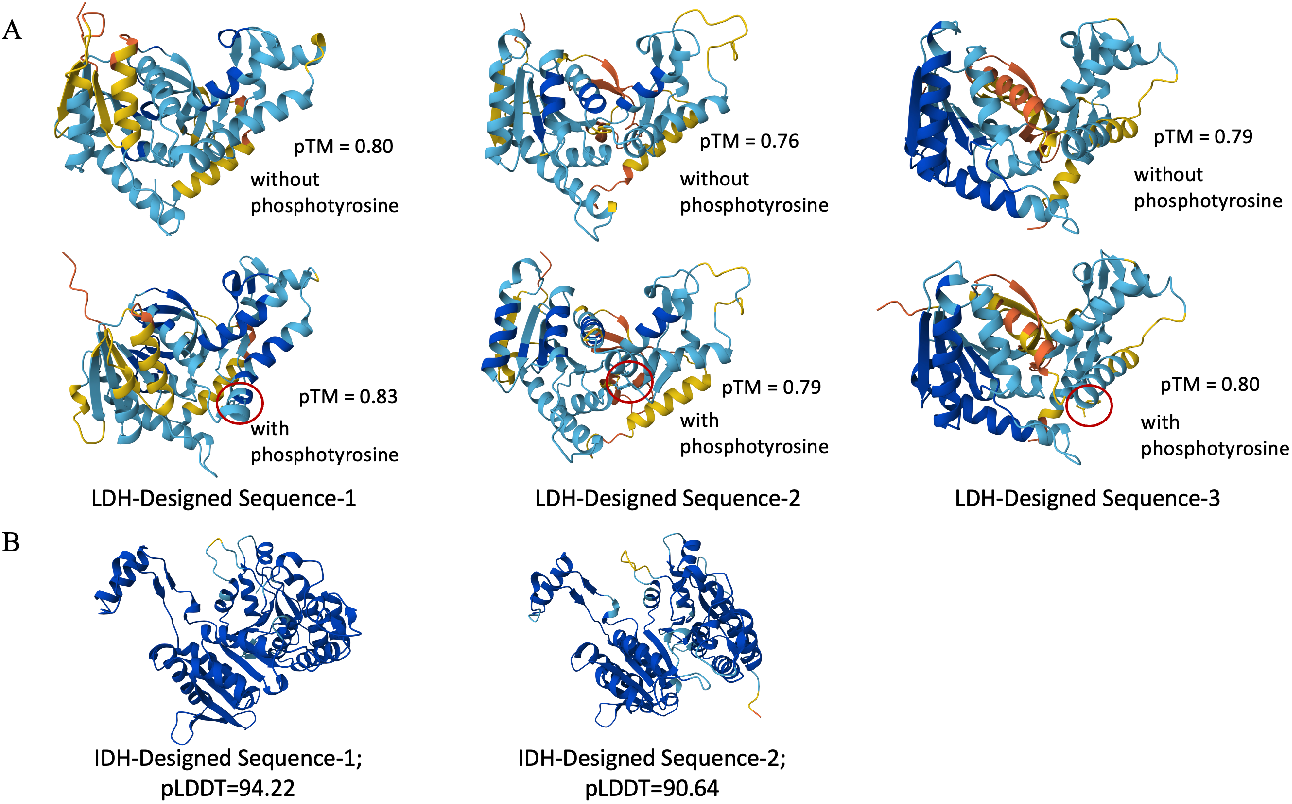
Predicted Structures of Unconditional Designed LDHs and IDHs.

### 2.3 Single-Site PTM Design of Lactate Dehydrogenase

We then evaluated whether PTM-dCN can precisely regulate the occurrence probability of an existing PTM site in a native protein sequence, a property that is closely related to biological processes such as molecular interaction and signal transduction.

Given that the occurrence of PTMs depends not only on the modified residue itself but also on the surrounding amino acids and adjacent chemical environment, we redesigned the sequence context of an existing phosphorylation site while preserving the remainder of the sequence.

We specifically target the Tyr233 phosphorylation site commonly annotated in LDH dataset. The 10 upstream and 10 downstream residues surrounding the modified tyrosine were masked, while the modified residue and the rest of the sequence remained unchanged. Then PTM-dCN was used to redesign the local sequence context around the modification site (Figure A.2). From this process, we generated 64 PTM-containing LDH sequences.

The generated sequences were subsequently evaluated using the PTMGPT2 web server Shrestha et al. (2024). Among the 64 redesigned sequences, 56 were predicted to retain Tyr223 phosphorylation, and all of these sequences exhibited a high degree of conservation in their phosphorylation motifs (Figure A.3 (A)). These results indicate that PTM-dCN is able to generate sequence contexts that are highly compatible with known PTM recognition patterns.

To further investigate whether PTM-dCN designs site-specific PTM through extrapolating global context information rather than introducing a fixed modification motif, we analyzed the contextual composition patterns of the redesigned phosphorylation sites across different LDH starting sequences.

Despite targeting the same Tyr223 phosphorylation site, the redesigned motifs displayed substantial diversity across different sequence backbones, suggesting that PTM-dCN adapts PTM-associated motifs in a backbone-dependent manner (Figure A.3). This behavior contrasts with traditional rational design strategies, which typically introduce predefined consensus motifs without learning the broader context within the whole sequence.

Having established that PTM-dCN can redesign existing PTM sites, we explored whether the model can upregulate the occurrence probability of low-confidence modification sites. Using MusiteDeep Wang et al. (2020), we screened LDH sequences containing a single annotated tyrosine phosphorylation site between residues 210–250 and with predicted phosphorylation probabilities below 0.7, resulting in 19 low PTM probability sequences. PTM-dCN was applied to redesign the context surrounding phosphorylation sites in these sequences. Remarkably, for each wild-type sequence, PTM-dCN generated redesigned variants exhibiting uniformly increased predicted phosphorylation probabilities, most of which exceeded 0.9 (Figure 4). These results demonstrate that PTM-dCN is capable of not only preserving but also enhancing the occurrence probability of existing PTM sites through context-aware sequence redesign.

**Figure 4:**
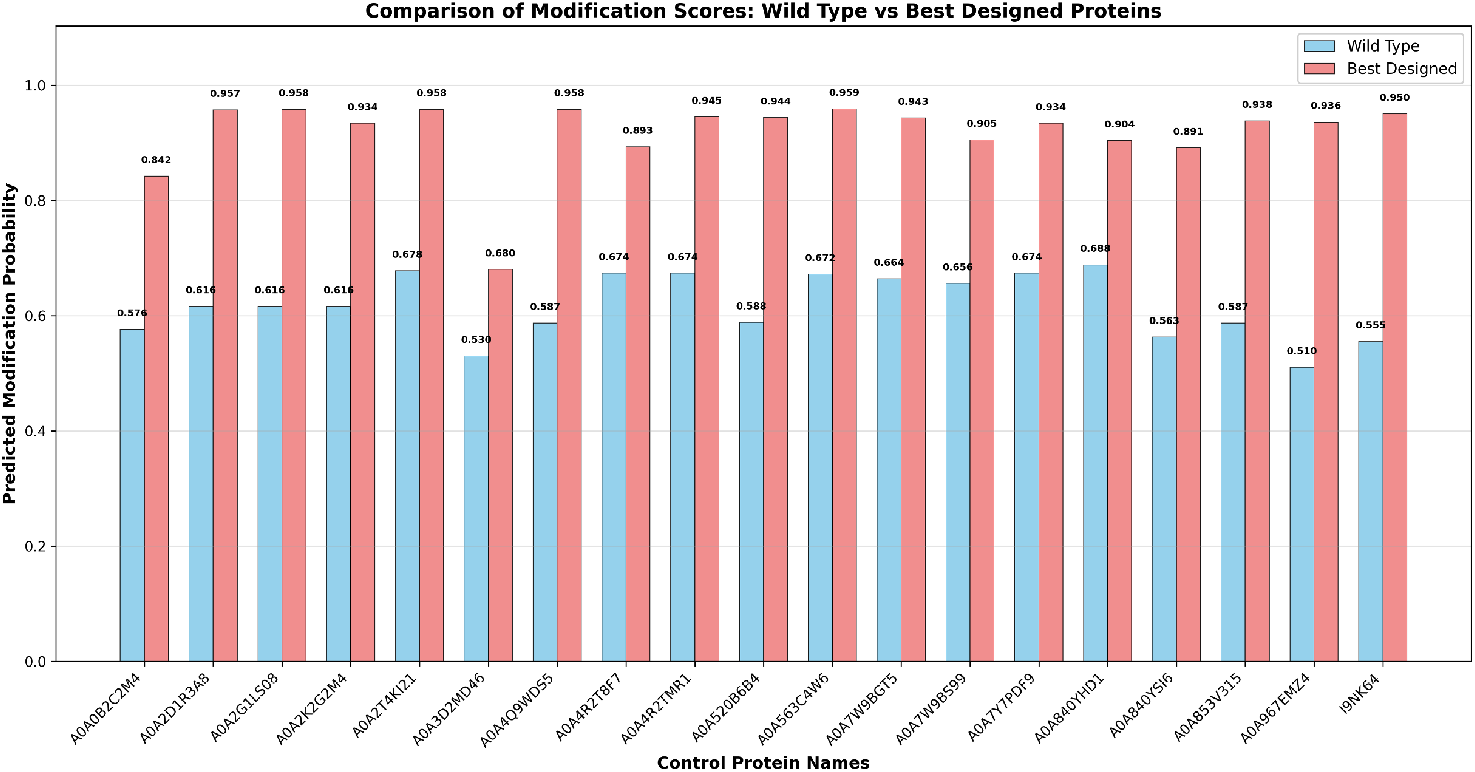
Predicted modification probability of wild type and best designed sequences.

### 2.4 Multiple-Site PTM Design of Isocitrate Dehydrogenase

Having demonstrated precise control over single-site PTM redesign, we further deployed PTM-dCN in the more challenging task of redesigning protein sequences containing multiple PTM types and sites. Based on a statistical analysis of PTM annotations (Figure A.4), isocitrate dehydrogenase (IDH; EC 1.1.1.42) was selected as a examplary system, which often contains three distinct PTM types: N6-succinyllysine (45.5%), phosphoserine (30.0%), and N6-acetyllysine (24.4%). In addition, 89.3% of IDH sequences fall within a length range of 301–500 amino acids, making this dataset suitable for controlled multi-site PTM design.

Following the procedures used for single PTM design in LDH, we trained both the diffusion backbone and the ControlNet module on the IDH dataset. When simultaneously conditioning the model on three distinct PTM types, we observed that the ControlNet module exerted weaker constraints on the generation process. Among 64 generated sequences, fewer than three matched the target multi-site PTM pattern (Figure A.5). Based on this observation, we hypothesize that, in the presence of higher sequence diversity and more complex PTM combinations, effective control requires a smoother latent space that is well-structured for Gaussian diffusion modeling.

To further investigate this phenomenon, we visualized the latent representations of both training and generated sequences using two-dimensional projections and channel-wise heatmaps. The resulting distributions revealed that generated sequences were clustered within a relatively compact region of the latent space (Figure A.6 (A)), accompanied by large inter-channel variance(Figure A.6 (B)), which may hinder efficient diffusion learning. Inspired by prior works such as CHEAP Lu et al. (2025b) and PLAID Lu et al. (2025a), we introduced a tanh-based nonlinearity function to smooth the latent space, after which the latent representations became more evenly distributed (Figure 5 (A)) with reduced channel-wise fluctuations(Figure 5 (B)).

**Figure 5:**
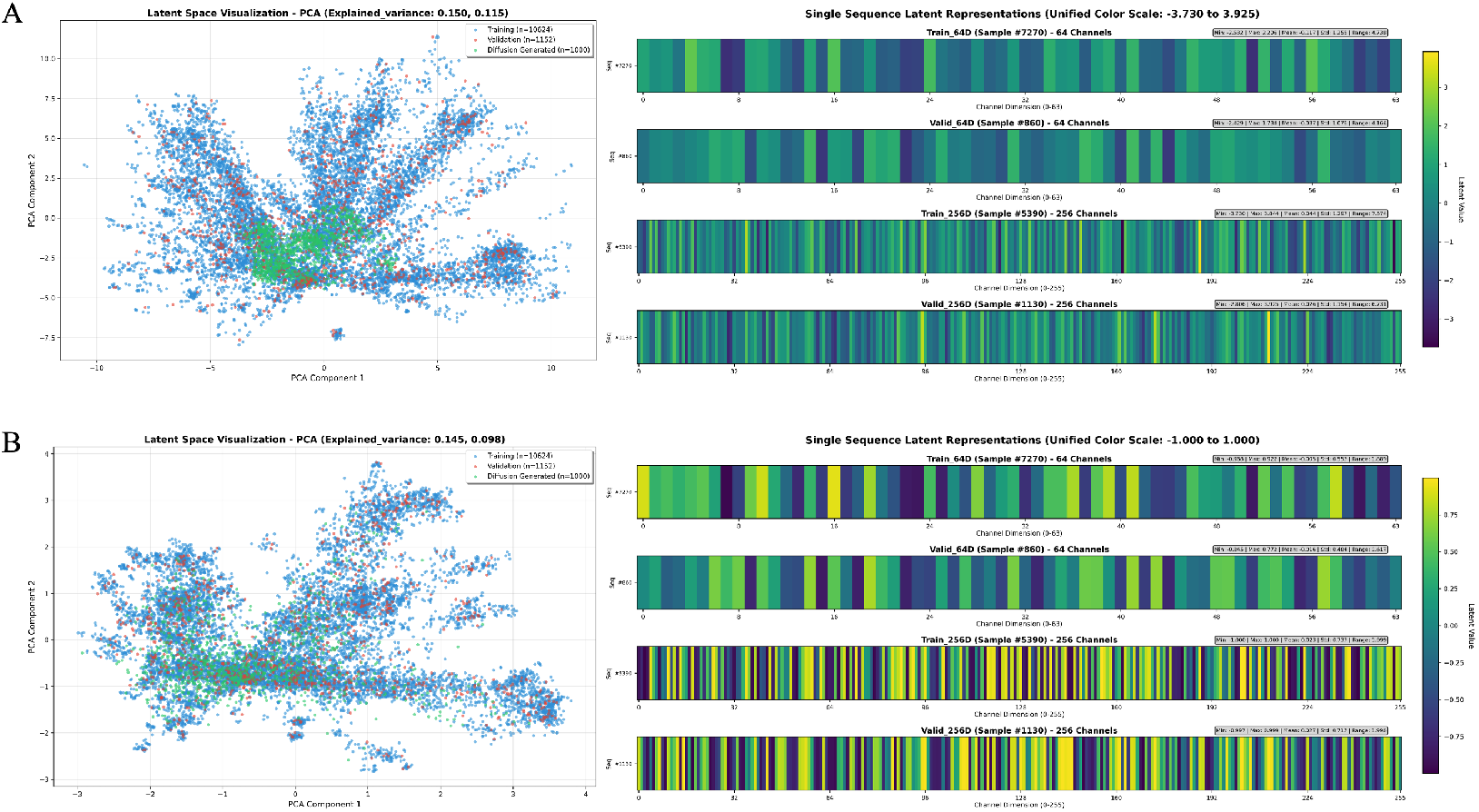
Low-Dimensional Visualization (left) and Heatmap (right) Analysis of Latent Representations Before (A) and After (B) Tanh Smoothing.

Following latent space smoothing and subsequent ControlNet training, the number of generated sequences matching the target multi-site PTM patterns increased substantially, from at most three to twelve sequences per 64 generations (Figure A.7). Notably, these sequences also exhibited high predicted foldability with typical pLDDT *>*70, indicating that latent space regularization plays a critical role in enabling effective and controllable multi-site PTM design.

## 3 Conclusion

In this work, we present PTM-dCN, a diffusion-based protein sequence generation framework that enables explicit, site-specific design of PTM patterns. By constructing a high-quality SwissProt-derived PTM-annotated dataset, introducing specialized PTM tokens, and integrating ControlNet into a latent diffusion backbone, our approach bridges global sequence modeling with precise engineering of local PTM sites and types.

Through comprehensive experiments on lactate dehydrogenase and isocitrate dehydrogenase, we demonstrate that PTM-dCN can (i) generate novel PTM-containing protein sequences with high predicted foldability, (ii) redesign local sequence contexts to preserve or substantially enhance PTM occurrence probabilities at target sites, and (iii) adapt PTM-associated motifs in a context-dependent manner rather than relying on fixed consensus patterns. Importantly, we show that latent space smoothness is critical for effective multi-site and multi-type PTM insertion, and that simple tanh-based regularization significantly improves controllability and design quality.

Overall, PTM-dCN provides a general and extensible framework for precise protein design with site-aware information, offering a new avenue for systematically exploring PTM-specific functional landscapes and advancing controllable protein engineering with deep generative models.

## Appendix

### A.1 Data and code availability

The model and training weights will be released on GitHub. Further information is available upon request from the corresponding author.

#### A.2 Dataset Construction

##### A.2.1 SwissProt PTM dataset

The SwissProt dataset (UniProt, May 2024 release) was collected along with corresponding “modified residue” annotations. Among 571,283 SwissProt entries, 74,671 protein sequences contain at least one annotated PTM, covering 354 distinct PTM types. The average sequence length of these sequences is 506 residues, and the PTM type distribution exhibits a long-tailed pattern, with 95 PTM types occurring more than 100 times (Figure A.8). Based on this criterion, 73,024 sequences containing PTM types with frequencies greater than 100 were selected to construct the SwissProt PTM dataset, which was randomly split into training and validation sets at a 9:1 ratio. During data preprocessing, the sequence length was truncated to 512 residues. For sequences longer than 512 residues, a sliding window strategy was applied to extract the subsequence containing the maximum number of modified residues, whereas sequences shorter than 512 residues were padded to a fixed length of 512 using a padding token.

##### A.2.3 LDH dataset

All protein sequences annotated with EC number 1.1.1.27 were retrieved from the UniProt database, including both SwissProt and TrEMBL entries. Sequence length and modified residue annotations were analyzed for this collection. Among the sequences containing annotated modified residues (7,232 in total), 98.9% (7,149 sequences) exhibit a single-site tyrosine phosphorylation located between residues 200 and 250 (Figure A.9). Based on this observation, these 7,149 sequences were retained to construct the LDH dataset, which was randomly split into training and test sets at a 9:1 ratio.

##### A.3 IDH dataset

All protein sequences annotated with EC number 1.1.1.42 were retrieved from the UniProt database, including both SwissProt and TrEMBL entries, and their sequence lengths and modified residue annotations were analyzed. Among the 15,518 sequences containing annotated modified residues, 89.3% (13,853 sequences) have lengths between 301 and 500 residues (Figure A.4). Across these 13,853 sequences, a total of 45,401 modifications were observed, with N6-succinyllysine, phosphoserine, and N6-acetyllysine accounting for 20,658 (45.5%), 13,634 (30.0%), and 11,092 (24.4%) occurrences, respectively. Based on these statistics, a dataset comprising 11,831 sequences was constructed by selecting sequences of length 301–500 residues containing exclusively these three PTM types (each sequence may contain one or more of these modifications). The resulting dataset was randomly split into training and validation sets at a 9:1 ratio.

##### A.3 Supplementary Figures

**Figure A.1:**
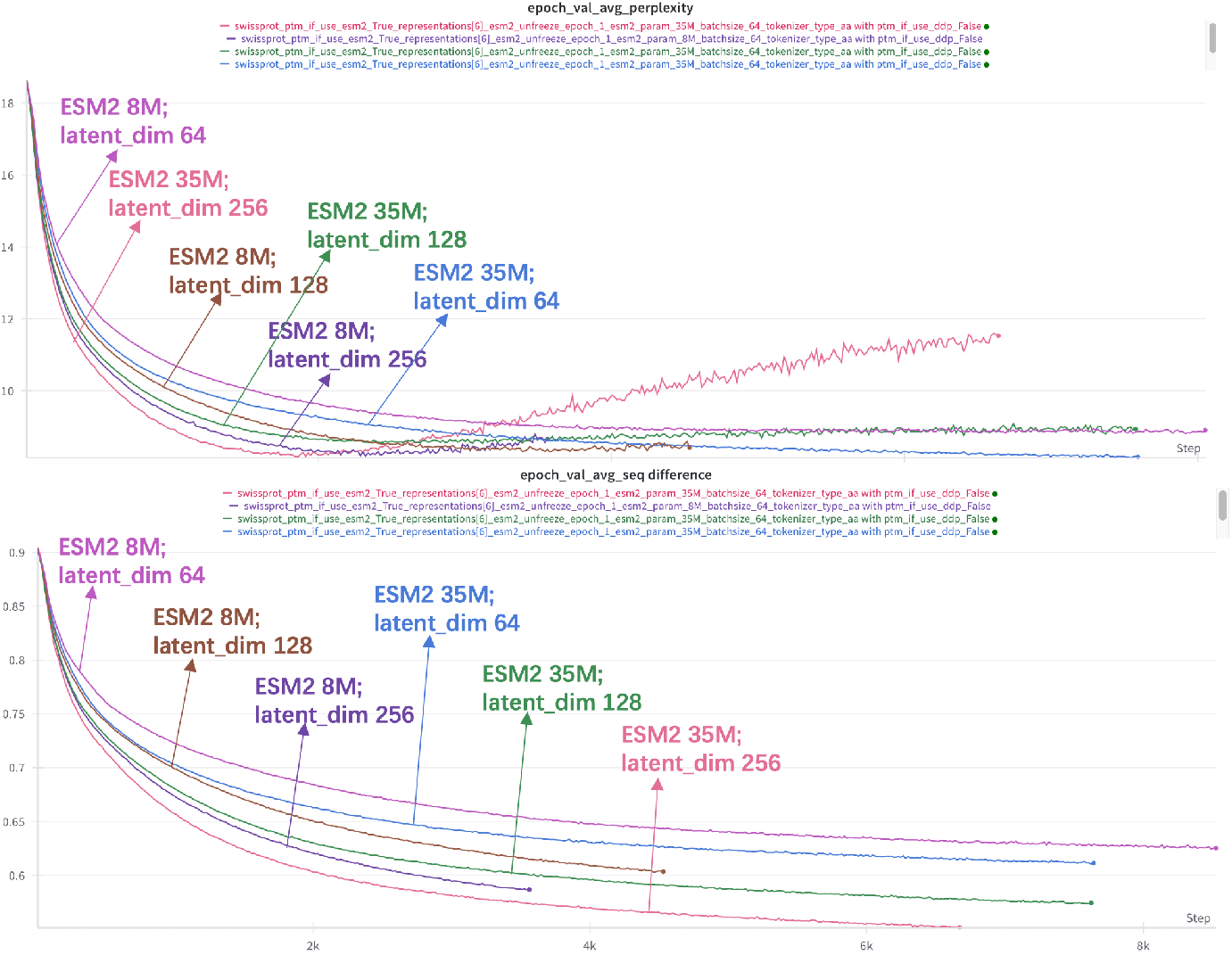
Evolution of Perplexity and Sequence Difference During Training.

**Figure A.2:**
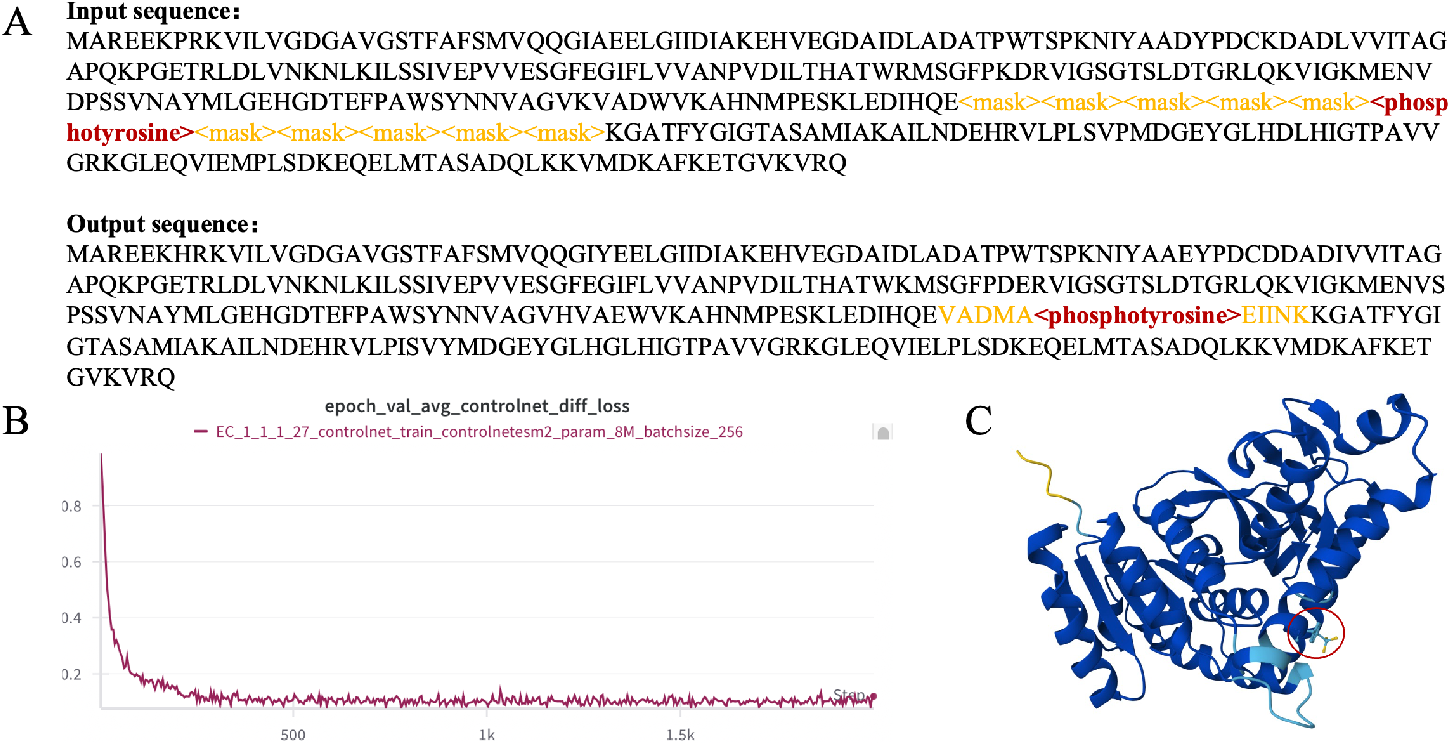
Generation Performance of PTM-dCN. (A) Examples of input sequence and output sequence. (B) Diffusion loss during training ControlNet module of validation set. (C) The AF3 predicted structure of redesigned LDH with phosphorylation site.

**Figure A.3:**
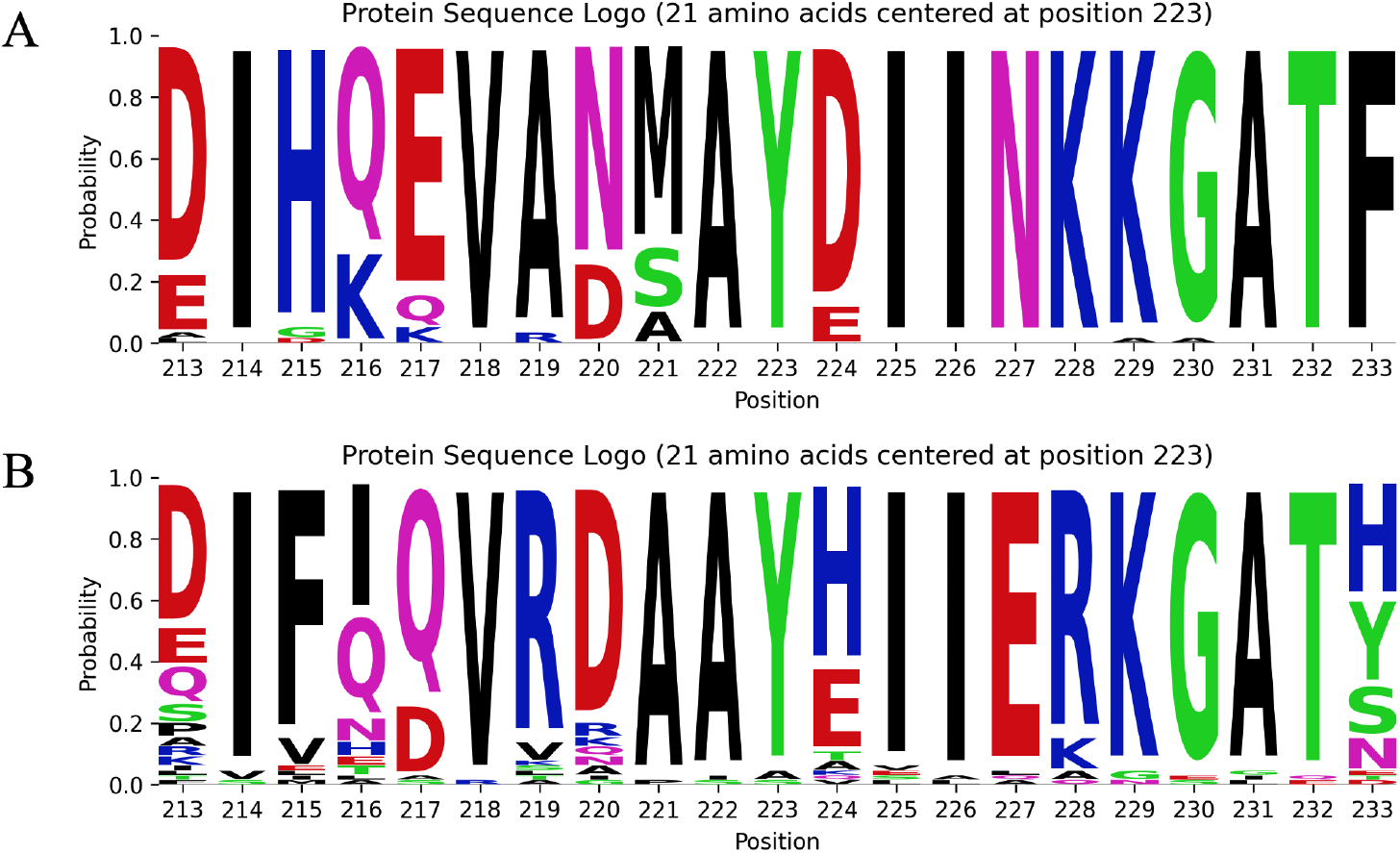
Logo figures of redesigned LDH sequences based on two different wild type seuquences. (A) Based on wild type LDH seuquence A; (A) Based on wild type LDH seuquence B.

**Figure A.4:**
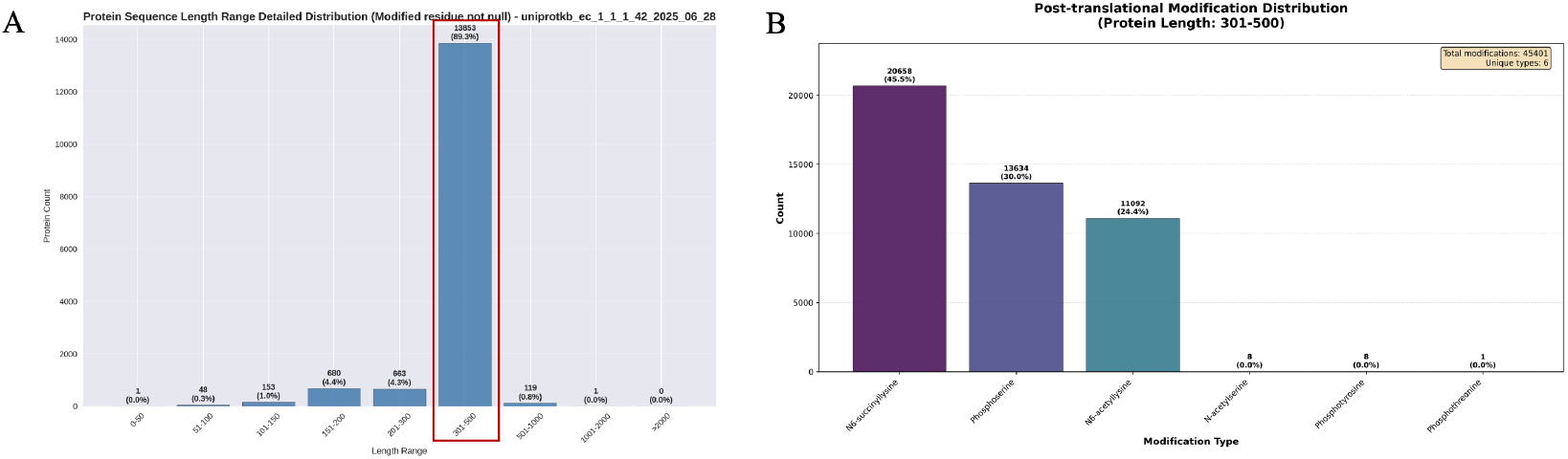
Protein sequence length (A) and PTM types (B) distribution of IDH dataset.

**Figure A.5:**
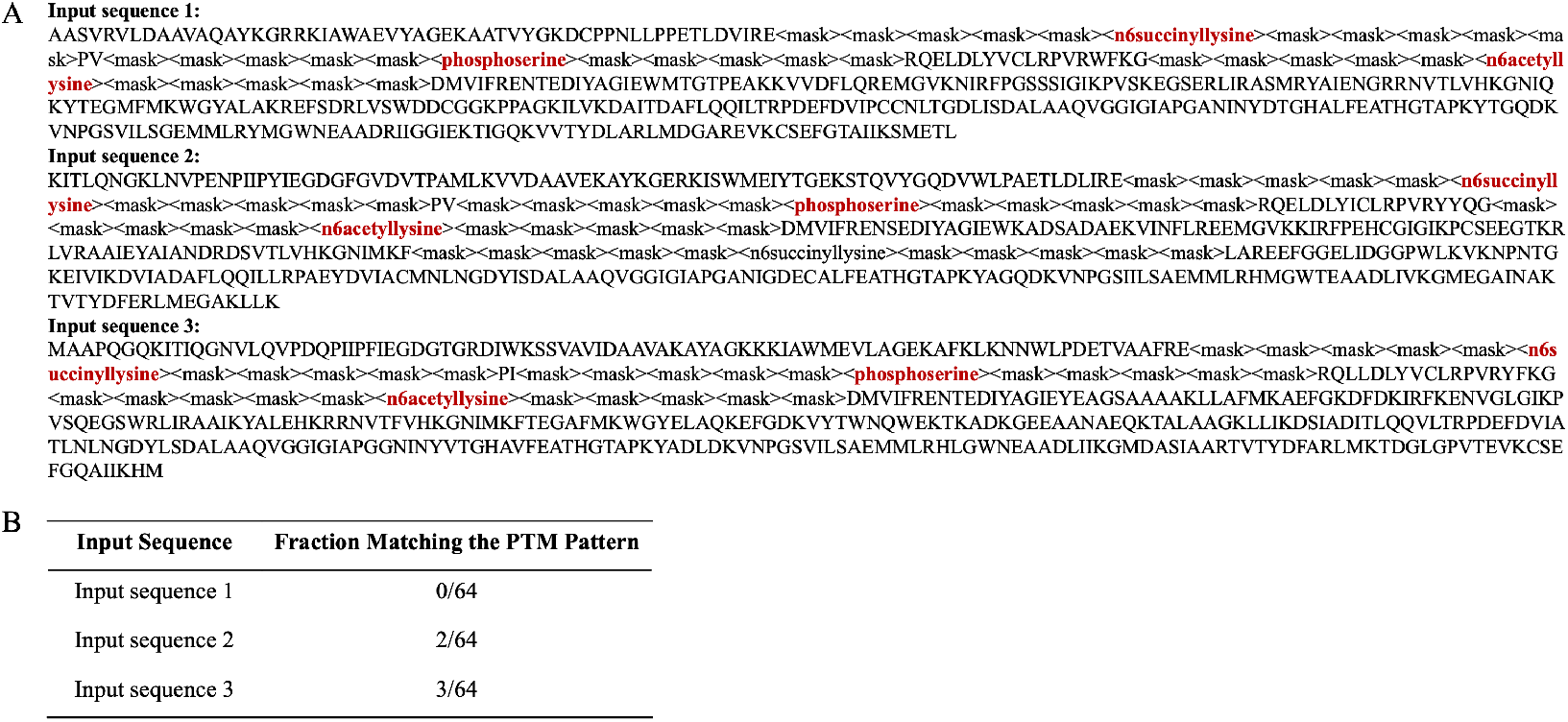
Input Sequences (A) and fraction matching the target PTM pattern (B).

**Figure A.6:**
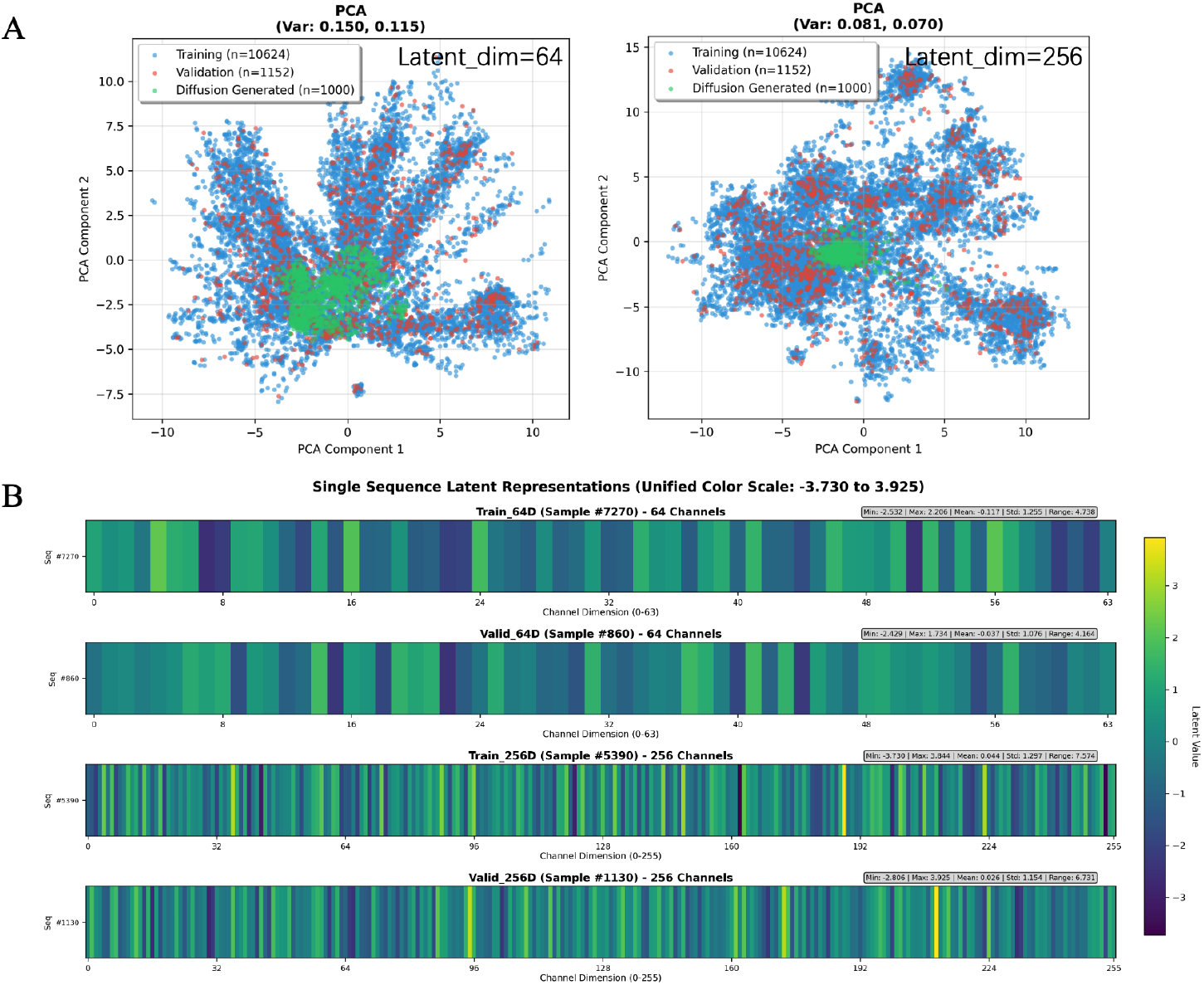
The embedding visualization of IDH sequences with different dimensions. (A) latent representation visualization comparision; (B) Heatmap of single sequence latent represention.

**Figure A.7:**
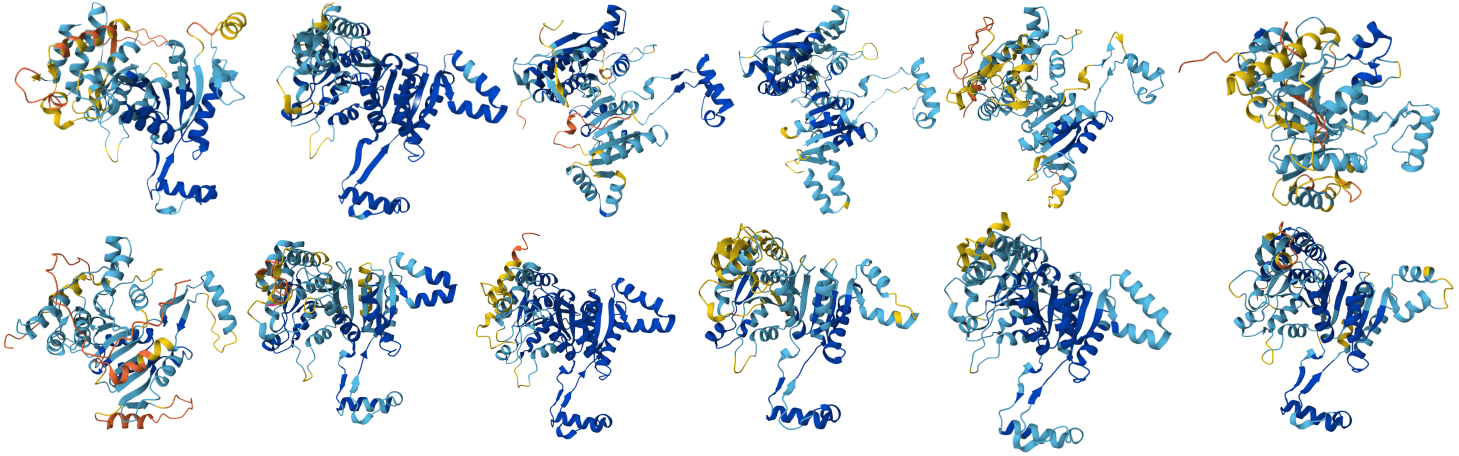
The predicted structures of 12 IDH generated sequences with target PTM tyeps.

**Figure A.8:**
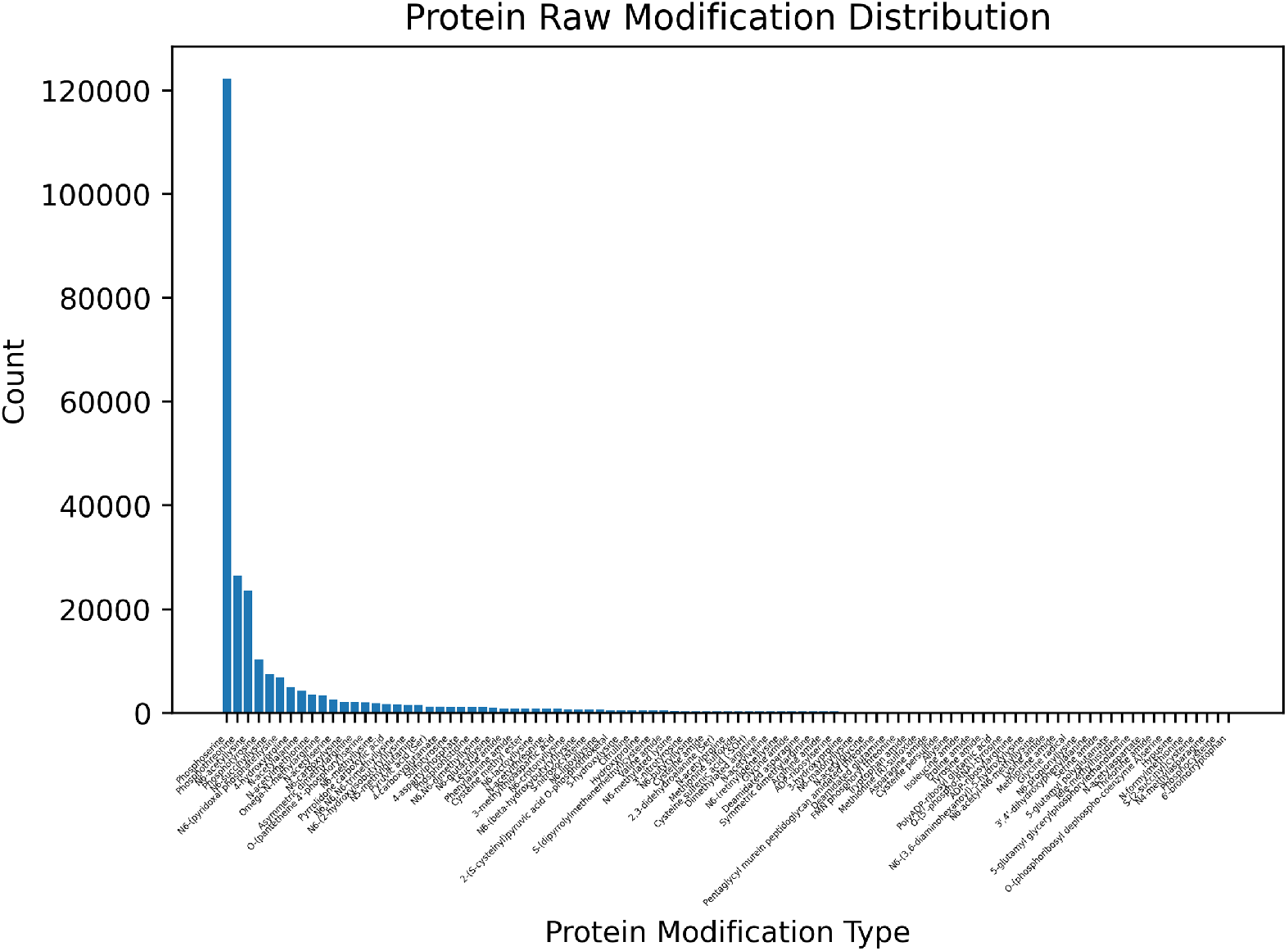
The distribution of all 354 PTM tyeps in SwissProt dataset.

**Figure A.9:**
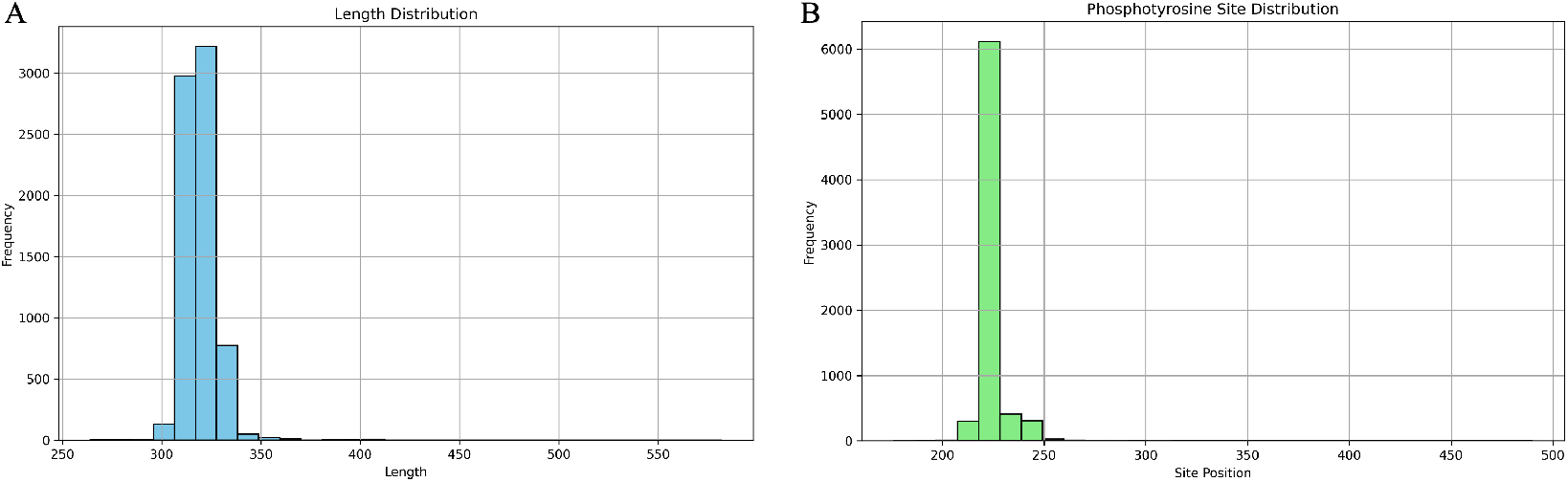
The distribution of length (A) and phosphotyrosine (B) in LDH dataset.

